# Transcriptional condensates formed by phase-separated ALOG family proteins control shoot meristem maturation for flowering

**DOI:** 10.1101/2021.03.18.435998

**Authors:** Xiaozhen Huang, Nan Xiao, Yue Xie, Lingli Tang, Yueqin Zhang, Yuan Yu, Cao Xu

## Abstract

Plants have evolved remarkable diversity in inflorescence architecture. At the center of this diversity lies a meristem maturation program featured by transition of stem cell populations from a vegetative state into a reproductive growth, determining when, where, and how many flowers are produced on inflorescences. Here we identified a new meristem maturation regulator *TMF FAMILY MEMBER3* (*TFAM3*) that encodes an ALOG family transcription factor. Loss of *TFAM3* results in early flowering and simplified inflorescences with fewer flowers. Genetic analysis by creating high-order mutants of *TFAM3* with three key regulators of tomato shoot meristem maturation, *TERMINATING FLOWER* (*TMF*), *TMF FAMILY MEMBER1* (*TFAM1*) and *TMF FAMILY MEMBER2* (*TFAM2*), suggested that they synergistically control flowering transition and inflorescence architecture. The four paralogous ALOG proteins share the prion-like properties and undergo liquid-liquid phase separation *in vitro*. Strikingly, TMF can recognize cognate TFAM proteins and selectively recruit them into phase separated condensates. Supporting this, they interact with themselves and each other to form biomolecular condensates in the nucleus. Their interaction induces formation of transcriptional condensates that directly repress expression of floral identity gene *ANANTHA*. Our study revealed a selective-recruitment phase separation mechanism for transcriptional condensation by which plants achieve optimal coordination of functional overlapped paralogs within a protein family to enable precise control of shoot meristem maturation for flowering and production of compound inflorescences.

## Introduction

Reproductive success and shoot architectural diversity of flowering plants rely on the time of flowering and the architecture of inflorescences (Benlloch et al., 2007; Rickett, 1944; Weberling, 1989; Xu et al., 2016). Inflorescences develop from shoot apical meristems (SAMs), whose fate are determined by balancing cell proliferation for replenishing pools of pluripotent cells with cell differentiation for organogenesis (Huang et al., 2021). Upon perception and integration of endogenous and environmental signals, SAMs experience a gradual maturation process from a vegetative stage (vegetative meristem, VM) into a floral stage (floral meristem, FM) accompanied by successive leaf production, called meristem maturation (Park et al., 2012, 2014).

The programed and successive meristem maturation ensures the transition to flowering at proper time and the formation of flowers in appropriate amount, which directly influence reproductive success of plants and yield of crops (Park et al., 2014; Xu et al., 2016). Before reaching the transition stage, SAM at vegetative stage is too juvenile to respond to endogenous or environmental signals. A program that prevents precocious maturation is vital for enabling a sufficient duration of vegetative stage for accumulation of adequate stem cells. Upon “juvenile” meristem maturing to “adult” meristem, a promoting program is required to activate floral identity gene for flowering. Therefore, both repressing and promoting programs during meristem maturation are essential for precise control of flowering transition and inflorescence architecture.

Over the past decades, studies in model and crop plants have reported genetic pathways that implicate in flowering transition and the formation of diverse inflorescence architectures (Barton, 2010; González-Suárez et al., 2020; Teo et al., 2014; Wu et al., 2018; Zhu and Wagner, 2020). Studies on tomato inflorescence mutants and wild species have come up with a working model that illustrates the mechanism for promoting meristem maturation. Once the promoting program is disrupted, meristem maturation is delayed or fails to complete, giving rising to over-proliferated axillary inflorescence meristems (AIMs) and thus highly branched inflorescences (Park et al., 2012). Loss-of-function of homeobox gene *COMPOUND INFLORESCENCE* (S) in tomato or *EVERGREEN* (*EVG*) in petunia, both genes are homologs of *Arabidopsis WOX9*, leads to highly branched inflorescences (Hendelman et al., 2021; Lippman et al., 2008; Rebocho et al., 2008). Mutant of the F-box gene *ANANTHA* (*AN*; homolog of *Arabidopsis UFO* and petunia *DOT*) in tomato never forms normal inflorescences and flowers, only produces overproliferated axillary inflorescence meristems (Hepworth et al., 2006; Lippman et al., 2008; Rebocho et al., 2008; Souer et al., 2008). Genetic studies together with tissue-specific transcriptome profiling of SAMs in tomato cultivar and wild species support a model wherein programmed activation of *S* followed by *AN* drives successive stages of axillary inflorescence meristems maturation to ensure them develop at precise time and in appropriate amount in each sympodial growth cycle (Park et al., 2012, 2014).

A repressing program that maintains meristem at a vegetative state is defined by a tomato ALOG (*Arabidopsis* LSH1 and *Oryza* G1) transcription factor, TERMINATING FLOWER (TMF) (MacAlister et al., 2012). TMF harbors a conserved ALOG domain featured by a putative DNA-binding domain derived from the XerC/D-like recombinases of a novel class of retrotransposons (Iyer and Aravind, 2012; Xu et al., 2016). Loss of *TMF* in tomato leads to much faster flowering and conversion of the multiple-flowered primary inflorescence into a single flower. These effects caused by precocious activation of *AN* due to loss of TMF’s repression, making a meristem mostly at a vegetative stage acquire the floral identity (MacAlister et al., 2012). Surprisingly, inflorescences that develop from side shoots of *tmf* mutant are normal, implying that *TMF* paralogs in ALOG family or other genes might synergistically act with *TMF* to regulate the inflorescence architecture. Indeed, the tomato ALOG family includes twelve members, which are named as TMF FAMILY MEMBERs (TFAMs) (Xu et al., 2016). Two additional ALOG family members, TFAM1 and TFAM2, form transcriptional complex with TMF via homo- and hetero-interactions to act together in preventing precocious meristem maturation (Huang et al., 2018). Studies in *Arabidopsis* and other nightshades (Solanaceae) have reported important roles of ALOG genes in floral organ specification, light signaling (Chen et al., 2019; Cho and Zambryski, 2011; Takeda et al., 2011; Zhao et al., 2004). In rice and wheat, several ALOG family members have been found to regulate lemma and hull specification, spikelet development, panicle branching and grain size (Nan et al., 2018; Sato et al., 2014; Yoshida et al., 2009, 2013). Notably, loss of *Marchantia polymorpha LATERAL ORGAN SUPRESSOR1* (*MpLOS1*), a *TMF* ortholog in the liverwort, causes mis-specified identity of lateral organogenesis and defects in apical meristem maintenance, suggesting its essential role in convergent evolution of lateral organogenesis (Naramoto et al., 2019). ALOG proteins represent a new plant specific transcription factor family with vital functions in meristem activity and reproductive development, however, the molecular mechanism of their transcriptional regulation is poorly understood.

The plant meristem maturation is a programmed developmental event driven by precise switch of gene repression and activation, which might require integration and transmission of endogenous and environmental signals into gene expression to be finely controlled in a relatively undisturbed micro-environment. Compartmentalization within a cell is a widely used strategy for complicated cellular processes and biological reactions in multicellular organisms. Besides canonical membrane-bound organelles, biomolecular condensates, the microscale membraneless compartments formed by liquid-liquid phase separation, have been recently found to spatially concentrate proteins and nucleic acids (Banani et al., 2017). To date biomolecular condensates formed by protein phase separation have recently been demonstrated as a new strategy for many cellular functions in yeast and animal systems, including cell polarity establishment and maintenance, cell signaling, cell and organ development, cell survival, and aging (Alberti and Hyman, 2021; Banani et al., 2017; Boeynaems et al., 2018; Strzyz, 2019; Wu et al., 2020). However, the mechanism of protein phase separation and the function of derived biomolecular condensates in plants are still largely uninvestigated (Emenecker et al., 2021; Fang et al., 2019; Huang et al., 2021; Zhang et al., 2020). In this study, we found that phase-separation induced formation of a transcriptional condensate times floral identity gene expression to program shoot apical meristem maturation in tomato.

## Results

### *TFAM3* is a new regulator of flowering transition and inflorescence architecture

Biological robustness is the ability of a system to maintain its function despite environmental or genetic perturbation (Diss et al., 2014). Genetic robustness can be achieved by functionally overlapped paralogs within a gene family. The phenomenon of this kind of redundancy is not rare, however, the molecular mechanisms linking the genotype to the phenotype are still poorly understood. The fact that TMF, TFAM1 and TFAM2 act synergistically in meristem maturation prompts us to systemically explore the roles of TMF family members in this process to uncover the mechanism underlying their paralogous interactions. Phylogeny analysis and protein alignment showed that twelve TFAM family members can be divided into two clades. Seven TFAMs, TMF and TFAM 1, 2, 3, 5, 9, 11, were clustered into one group, in which TFAM3 is the closest paralog of TMF (Figure 1A). To focus on the potential partners of TMF and TFAM1/2, we first examine the expression pattern of TFAMs in meristems at various developmental stages. The tissue-specific transcriptome profiling data suggested that four members in this clade, *TMF* and *TFAM1, 2, 3, 11*, show predominantly expression in meristems (Park et al., 2012). Among them, *TFAM3* shows similarity to *TMF* and *TFAM1, 2*, which express highly during VM and TM stages and then decrease in FM, but *TFAM11* shows low expression in VMs and high expression in FM, being distinct from that of *TMF* (Figure 1B). In this regard, we use CRISPR/Cas9 system to knock out *TFAM3* (Figure 1C). The *tfam3* null mutant flowers about one leaf earlier and produces fewer flowers on the primary inflorescence compared with wild type (Figure 1D-F), resembling mutant phenotypes of *TMF* weak alleles (Huang et al., 2021; MacAlister et al., 2012). Moreover, about 13% of the primary inflorescences of *tfam3* show vegetative reversion, indicated by outgrowth of leaves on the inflorescence (Figure 1D). The *tfam3* mutant frequently underwent a single branching event on each inflorescence (Figure 1D). These phenotypes remind us the inflorescence development defects of *tfam1* and *tfam2* mutants. As previously reported and shown here, the flowering time is unaffected in *tfam1* and *tfam2* single mutants (Figure 1D and 1E). Instead, *tfam1* shows reduced flower production but highly frequent vegetative reversion on inflorescences, and *tfam2* develops inflorescences with a high frequency of single branching event (Figure 1D and 1F) (Xu et al., 2016). These findings suggest that *TFAM3* is a novel regulator that represses flowering transition and promotes inflorescence complexity. The partially phenotypic similarity of the three *tfam* mutants suggests their functional overlap.

**Figure 1.**
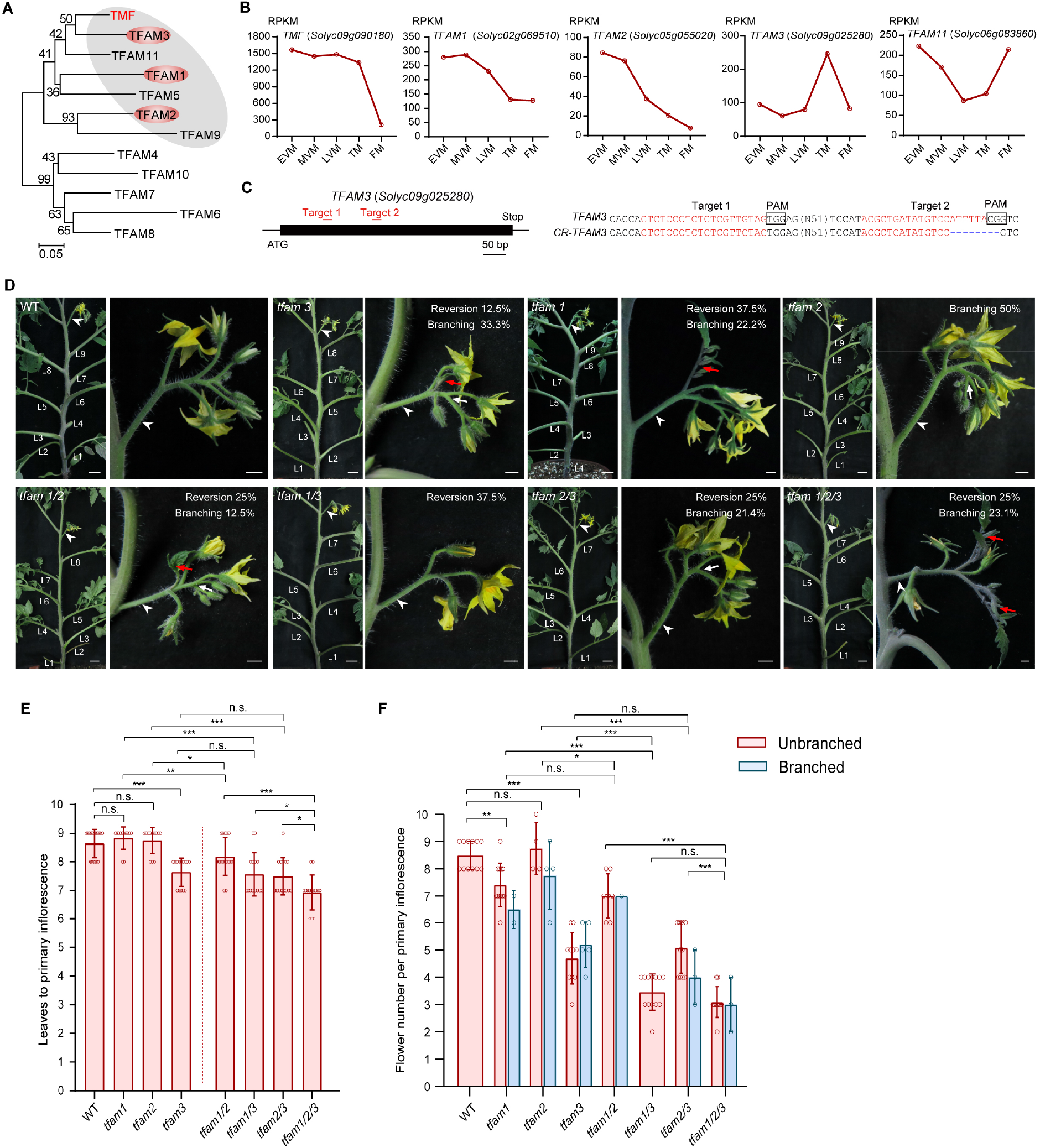
Identification of TFAM3 as a new regulator of flowering transition. (**A**) Phylogenetic tree showing the evolutionary relationship of ALOG family members in tomato. Red oval marked genes showing similar expression dynamics to *TMF* in meristems. (**B**) Expression dynamics for normalized read counts of *TMF*, *TFAM1*, *TFAM2*, *TFAM3*, and *TFAM11* at indicated stages during meristem maturation. (**C**) Schematic (left) indicating two sgRNAs (red lines) and allelic information (right) for *TFAM3*. (**D**) Representative shoot and typical primary inflorescence from WT and *tfam* single, double and triple mutants. White arrowheads indicate inflorescences, red arrows indicate vegetative reversions, white arrows indicate branching events on inflorescences. L, leaf. Scale bars, 2 cm for plants; 0.5 cm for inflorescences. (**E and F**) Statistics of flowering time (**E**) and flower number per inflorescence (**F**) for WT and *tfam* single, double and triple mutants. The flower numbers were quantified from branched and unbranched inflorescences separately. Data are means ±SD (n=17, 12, 12, 14, 16, 14, 14, 14, for **E**; n=12, 9, 8, 15, 8, 13, 14, 13 for **F**, *P*<0.05, \*\**P*<0.01, \*\*\**P*<0.001, Student *t*-test).

### Higher-order *tfam* mutants show enhanced early-flowering and inflorescence defects

To explore how *TFAM3, TFAM1* and *TFAM2* coordinate in synchronizing flowering transition and promoting inflorescence complexity, we generated various combinations of higher-order mutants by genetic crosses. Loss of either *TFAM1* or *TFAM2* in *tfam3* mutant background enhances its early flowering phenotypes, showing one leaf earlier than *tfam3* single mutant, about two leaves earlier than wild type plants (Figure 1D and 1E). The *tfam1/2* flowered faster than either single mutant by about one leaf, showing weaker early-flowering phenotype than *tfam1/3* and *tfam2/3* (Figure 1D and 1E). In addition to enhanced flowering phenotypes, the *tfam* double mutants showed a range of modifications to inflorescence architecture. Most dramatically, *tfam1/3* inflorescence produces only less than half of the number of flowers that the wild-type plant does, and about 38% inflorescences show vegetative reversion, similar to that of *tfam1* but stronger than that of *tfam3* (Figure 1D–1F). In contrast, *tfam2/3* and *tfam1/2* only displayed a slight change of the frequency of vegetative reversion and branching in comparison with single mutants (Figure 1D–1F). Strikingly, comparing to *tfam* doubles, the *tfam1/2/3* triple mutant flowered even faster, and its inflorescences only produced about three flowers (Figure 1D–1F). The progressive enhancement of flowering and inflorescence defects displayed by a complete series of *tfam* mutants indicate that their inseparable relationship in modulating flowering time and inflorescence architecture.

### *TMF* acts together with *TFAMs* to control flowering and inflorescence complexity

The *TMF* gene is a core member of ALOG family in tomato and plays pivotal role in regulating flowering and inflorescence complexity (MacAlister et al., 2012; Xu et al., 2016). The early-flowering and simplified-inflorescence phenotypes of various *tfam* mutants are similar, albeit weaker, to *tmf* null mutant. We therefore hypothesize that TMF might function as a “leader” to recruit and organize three TFAMs. To address this, we crossed all the single and multiple *tfam* mutants with *tmf* to create a series of *tmf tfam* mutant combinations. Among the double mutants, *tmf tfam1* and *tmf tfam3* showed the most significant enhancement of early-flowering comparing to *tmf* (Figure 2A and 2B). In contrast, there is no significant difference in flowering between *tmf tfam2* and *tmf* (Figure 2A and 2B). More prominent enhancement of early-flowering occurs in triple mutants. The *tmf tfam1/2* and *tmf tfam1/3* flowered earlier than *tmf* single mutant by one and two leaves, respectively (Figure 2A and 2B). Strikingly, the *tmf tfam1/2/3* quadruple mutant flowered extremely early after producing only two leaves and developed single-flowered inflorescences (Figure 2A and 2B). In addition, the *tmf* single-flower phenotype showed about 80% penetrance in our growth condition, neither introduction of *TFAM1*, *2* mutation individually nor simultaneously significantly improves the penetrance. However, the *tmf tfam1/2/3* quadruple mutant showed almost 100% penetrance for the single-flower phenotype (Supplemental Figure 1). *tmf* is single-flowered, and this flower often develops leaf-like sepals (MacAlister et al., 2012; Xu et al., 2016). Interestingly, as introduction of more mutations of *TFAM* genes into *tmf* background, sepals of the solitary flower showed more leaf characteristics (Figure 2A). These findings suggest that *TMF* requires *TFAM*s to achieve the precise control of flowering transition, among which *TFAM1* and *TFAM3* contribute much more than *TFAM2*.

**Figure 2.**
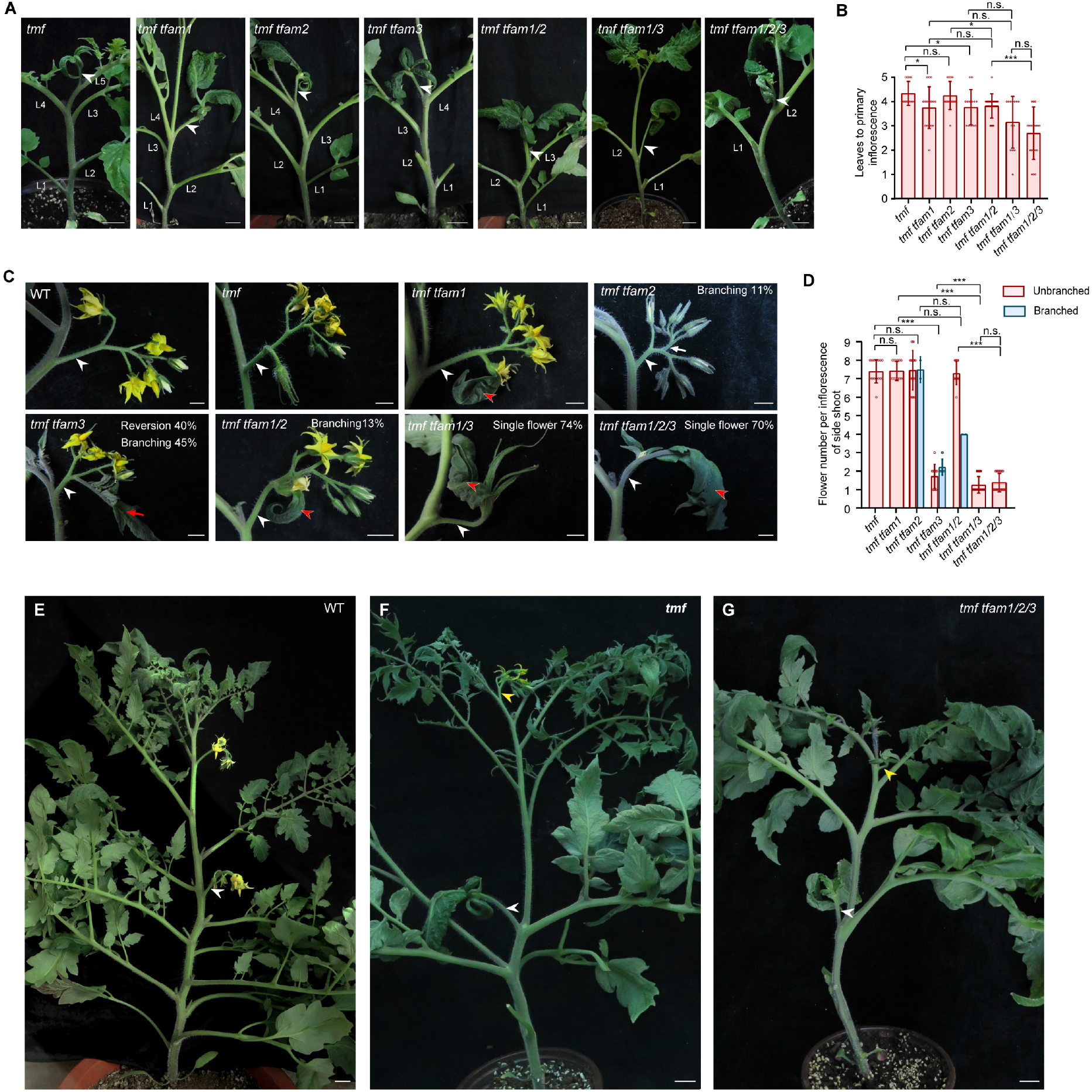
Genetic interactions between TMF and TFAM genes. (**A and B**) Representative shoots with primary inflorescence (**A**) and quantification of flowering time (**B**) for *tmf* single mutant and higher-order mutants of *tmf* and *tfams*. White arrowheads indicate single-flowered primary inflorescences. Data are means ±SD (n=12, 16, 16, 13, 22, 13, 20, \**P*<0.05, \*\**P*<0.01, \*\*\**P*<0.001, Student *t*-test). L, leaf. Scale bars, 2 cm. (**C and D**) Images of inflorescence (**C**) and quantification of flower number per inflorescence (**D**) from side shoots of various mutant combinations. Red arrowheads indicate leaf-like sepal, and white arrowheads indicate inflorescences. Data are means ±SD (n=15, 14, 19, 20, 15, 22, 25, \*\*\**p*<0.001, Student *t*-test). Scale bars, 1 cm. (**E-G**) Representative shoot with two successive inflorescences for WT (**E**), *tmf* single mutant (**F**) and *tmf tfam1/2/3* quadruple mutant (**G**). White arrowheads and yellow arrowheads indicate primary inflorescences and side-shoot inflorescences, respectively. Scale bars, 2 cm.

While the primary inflorescence of *tmf* is single-flowered, inflorescences that develop from side shoots are unaffected (MacAlister et al., 2012), suggesting the existence of redundant factors. We then examined the inflorescences from side shoots of various higher-order mutants combined by *tmf* and *tfams*. Quantification of flower number per inflorescence from side shoots showed no significant difference between *tmf*, *tmf tfam1*, *tmf tfam2*, *tmf tfam1/2* mutants and wild type plants (Figure 2C and 2D), however, *tmf tfam1* and *tmf tfam1/2* displayed leaf-like sepal at the first flower on the side shoot inflorescences (Figure 2C). Notably, the side shoot inflorescences of *tmf tfam3* double mutant almost always produce only two flowers, and most of the inflorescences showed vegetative reversion and branching (Figure 2C and 2D). Most remarkably, approximately 74% of side shoot inflorescences from *tmf tfam1/3* are single-flowered with extremely leaf-like sepals (Figure 2C and 2D), which never appears in *tfam1/2/3* triple mutants (Figure 1D and 1F). Interestingly, the side shoot inflorescences of *tmf tfam1/2/3* are mostly undistinguished from *tmf tfam1/3*, suggesting nonessential role of TFAM2 in this process (Figure 2C-2G). Together, these results suggest that *TMF*, *TFAM1* and *TFAM3* indispensably work together in flower production on both primary and side shoots.

### TMF and TFAMs synergistically repress shoot apical meristem maturation

To explore the developmental basis for the flowering and inflorescence defects in various single and high-order mutants, we dissected and compared the shoot apical meristems at reproductive stages. Shoot apical meristems of the *tfam* single and higher-mutants are indistinguishable at the transitional meristem (TM) stage (Figure 3A). However, the maturation rate of the SAM, indicated by the number of leaf primordium produced before vegetative meristems transitioning into floral meristems, varied in different mutants. Neither *tfam1* nor *tfam2* showed modified maturation rate, however, *tfam3*, *tfam1/2* and *tfam2/3* transitioned faster than wild type by about one leaf primordium (Figure 3A and 3B). *tfam1/3* and *tfam1/2/3* exhibited the fastest maturation rate, producing about two leaf primordium fewer than wild type for floral transition (Figure 3A and 3B). The faster maturation gave rise to the early-flowering phenotypes in those mutants. The inflorescence complexity can be reflected by the number of AIM initiated at young inflorescence stage (Xu et al., 2016). *tfam1/2* shows slightly slower initiation of the AIMs, however, *tfam1/3, tfam2/3 and tfam1/2/3* initiated remarkably fewer AIMs than wild type (Figure 3C). The precocious termination of the AIM initiation interpreted the simplified inflorescences of these *tfam* mutants. When examining the young inflorescence stages of various combinations of *tmf* and *tfam* mutants, one can clearly observe extremely simplified inflorescences featured by single flowers with leaf-like sepals (Figure 3D). The progressive enhancement of flowering and inflorescence defects were reflected by the number of leaf primordium production and the size of leaf-like sepals (Figure 3D and 3E). Taken together, these results indicated that TMF and TFAMs synergistically prevent precocious maturation of shoot apical meristems to ensure appropriate flowering transition and inflorescence complexity.

**Figure 3.**
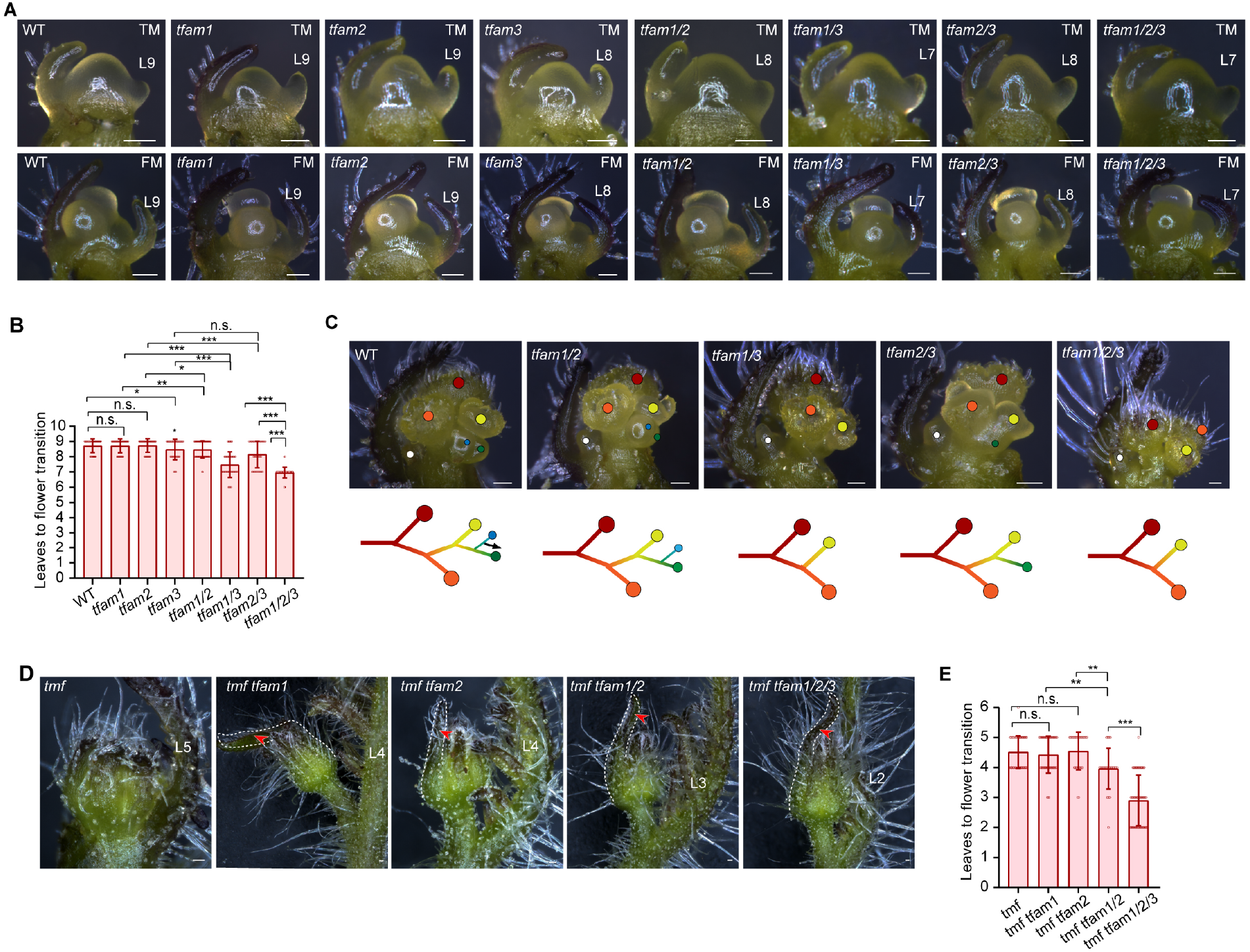
Developmental basis of shoot apical meristem maturation in *tmf* and *tfam* mutants. (**A and B**) Stereoscope images of meristems (**A**) and quantification data of leave primordium production for flower transition (**B**) from WT and *tfam* single, double and triple mutants. Data are means ±SD (n=69, 46, 46, 42, 59, 54, 39, 73, \**P*<0.05, \*\**P*<0.01, \*\*\**P*<0.001, Student *t*-test). Scale bars, 100 μm. L, leaf. (**C**) Young inflorescences (upper) and diagrams (bottom) of WT and *tfam* mutants. Colored dots indicate terminated FM (red, orange and yellow dots) and initiated SIM (blue and green dots). White dots indicate the first SYM. The black arrow indicates continued SIM reiteration. Scale bars, 100 μm. (**D and E**) Stereoscope images of floral meristem (**D**) and quantification of leaf production for flower transition (**E**) from *tmf* and *tfam* mutants. Data are means ±SD (n=53, 42, 29, 29, 62, \*\**P*<0.01, \*\*\**P*<0.001, Student *t*-test). Red arrowheads indicating the leaf sepal at floral meristem stage. Scale bars, 100 μm. L, leaf.

### TFAM proteins form biomolecular condensates in the nucleus of tomato cells

The synergistic interactions and overlapped functions prompt us to investigate if TMF and TFAM proteins share conserved properties. We analyzed the protein primary structure and found that all three TFAMs share highly conserved ALOG domain with a zinc-ribbon motif inserted in the middle region, which is a putative DNA-binding domain derived from the XerC/D-like recombinases. They also harbor a classic nuclear localization signal (NLS) in the C-terminus (Figure 4A). Several ALOG family proteins, including TMF, have been reported to have transcriptional activity (Huang et al., 2021; Iyer and Aravind, 2012; MacAlister et al., 2012; Yoshida et al., 2013). Transcriptional activity assays in yeast showed that TFAM2 and TFAM3 displayed significantly transcriptional auto-activation in the yeast-two hybrid systems, however, no auto-activation was detected for TFAM1, probably due to its unique DNA-binding preferences (Figure 4B). To explore the protein behavior of TFAMs in living tomato cells, we expressed TFAM-GFP fusion proteins in tomato protoplast. Confocal imaging showed that all three TFAM proteins exclusively localized in the nucleus as TMF did (Figure 4C). Strikingly, the GFP signals in the nucleus show high heterogeneity that marks an aggregate or condensate state, mimicking the punctate localization pattern of TMF (Figure 4D) (Huang et al., 2021). Given the fact that TMF undergoes liquid-liquid phase separation to form transcriptional condensates in the nucleus (Huang et al., 2021), the heterogenous condensation of TFAM proteins in the nucleus is likely due to protein phase separation.

**Figure 4.**
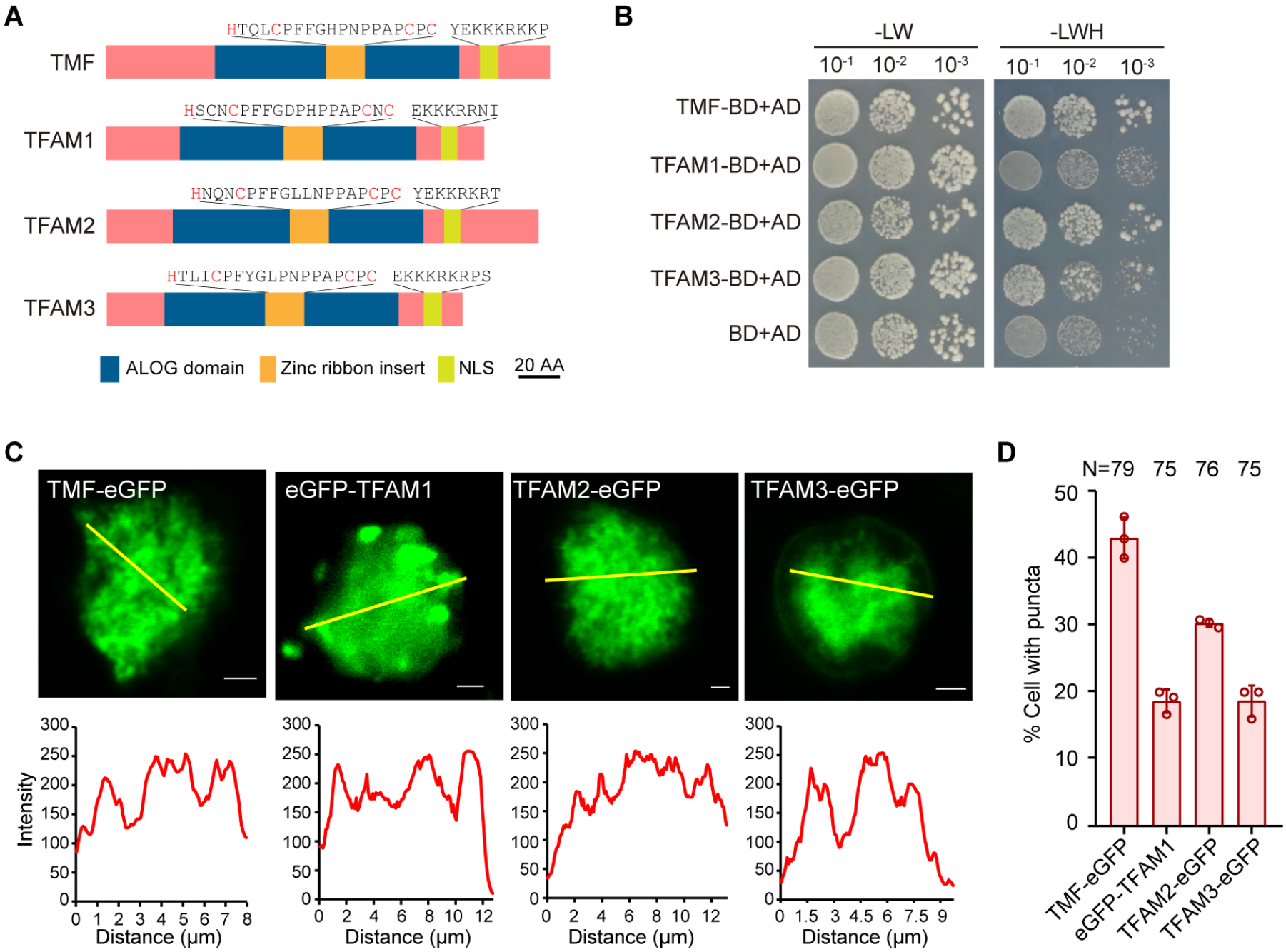
Protein behavior of TFAM proteins in yeast and tomato cells. (**A**) Schematics illustrating protein domains for TMF and TFAM proteins. (**B**) Yeast two-hybrid showing the auto-activation effects of TFAM proteins. Yeast cells were screened on the medium: SD-LW (-Leu/-Trp) and SD-LWH (-Leu/-Trp/-His). AD, activation domain; BD, DNA-binding domain. (**C**) Subcellular localization of TFAMs showing condensates in the nucleus of tomato cells (upper) and fluorescence intensity of indicated yellow lines (bottom). Scale bars, 2 μm. (**D**) Quantitative data showing the percentage of cells with condensates for GFP fusion proteins of TMF, TFAM1, TFAM2 and TFAM3 in the nucleus. Data are presented as three biological replicates ±SD (n=79, 75, 76, 75).

### TFAM proteins undergo phase separation *in vitro*

Further analysis of the TFAM proteins revealed that, like TMF, all three TFAMs have prion-like intrinsically disorder regions (IDRs) that are usually considered as a signature for protein phase separation (Figure 5A). We then recombinantly expressed and purified the TFAM-GFP fusion proteins from *E. coli* (Supplemental Figure 2A). We used the purified proteins to perform an *in vitro* phase separation assay that can generate a phase diagram by systematically changing protein and salt concentrations to assess the conditions that promote condensate formation (Huang et al., 2021). Interestingly, while all three TFAM proteins underwent phase separation, they showed some varying properties. Like TMF, TFAM3 readily phase-separated into droplets with a relatively regular spherical shape, however, TFAM1 and TFAM2 formed more irregular filamentous assemblies, these filaments and droplets are stable during the period of observation (Figure 5B-5D). The phase diagram showed a progressive increase of the density and size of the condensates formed by phase-separated TFAMs as the protein concentration improves when the salt concentration is constant (Figure 5B-5D). In contrast, the condensate abundance decreased with the increase of salt concentration when the protein concentration is constant, indicating they are sensitive to salt and protein concentrations (Figure 5B-5D). In particular, the TFAM3 proteins started to form visible spherical droplets at a concentration of 1 μM in a buffer with 150 mM NaCl (a physiologically relevant salt concentration), and the droplets rapidly fused together to form large droplet clusters as protein concentration increases (Figure 5D). It seems that the phase separation capacity of TFAM2 is weaker than that of TFAM1 and TFAM3 since it is more sensitive to salt.

**Figure 5.**
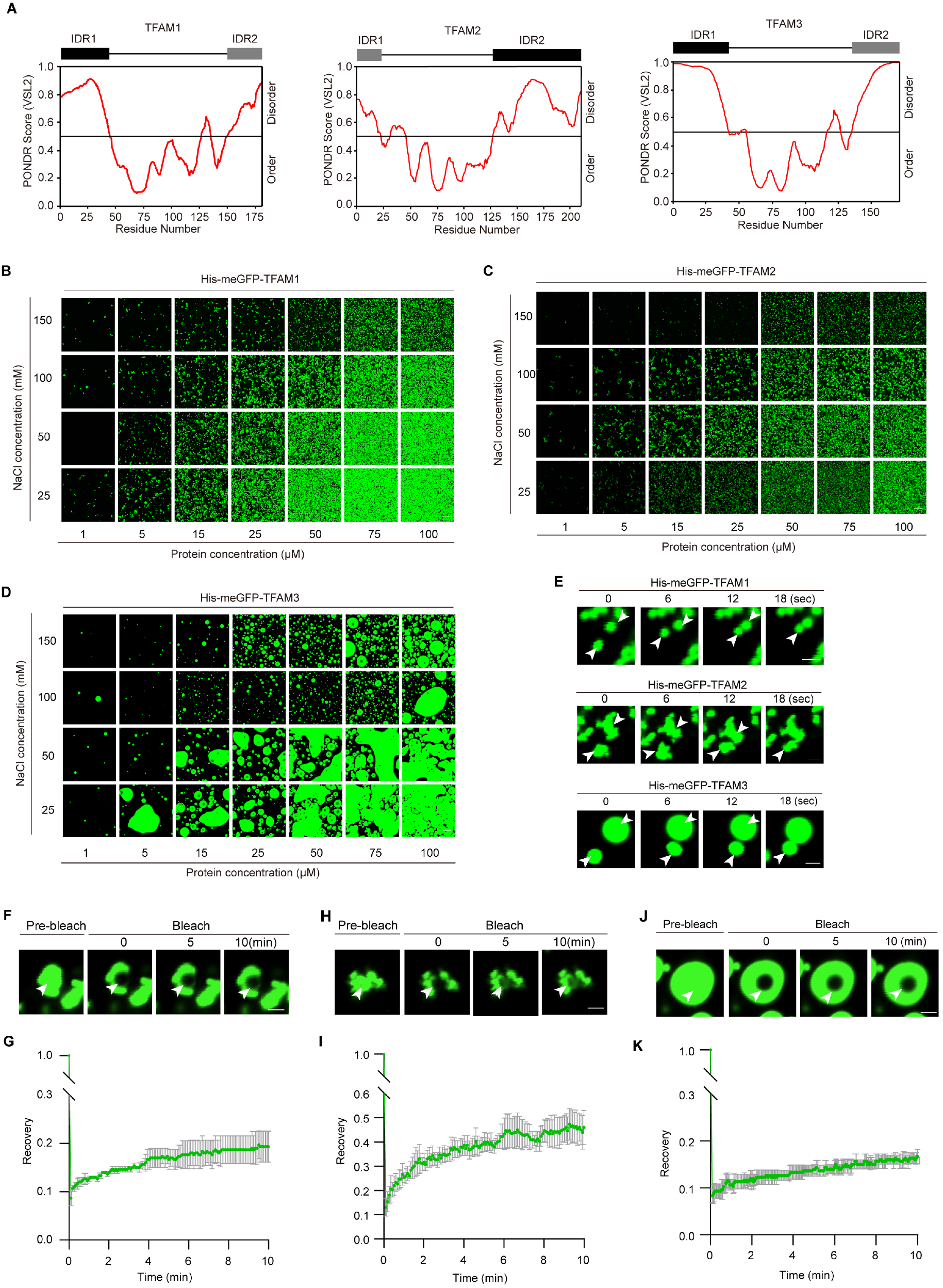
TFAM proteins undergo phase separation *in vitro*. (**A**) Graphs showing IDRs of TFAM proteins. (**B-D**) Phase separation of GFP-TFAM1 (**B**), GFP-TFAM2 (**C**) and GFP-TFAM3 (**D**) under the different combinations of indicated concentrations for NaCl and proteins. Scale bars, 20 μm. Three independent experiments with similar results were performed. (**E**) Representative images from three independent fusion events showing the liquidity of GFP-TFAM1 (upper), GFP-TFAM2 (middle) and GFP-TFAM3 (bottom) during phase separated droplets formation. Protein concentration, 15 μM; NaCl concentration, 25 mM. Scale bars, 2 μm. (**F-K**) Representative images and quantification data of FRAP analysis for GFP-TFAM1 (**F** and **G**), GFP-TFAM2 (**H** and **I**) and GFP-TFAM3 (**J** and **K**). Data are means of three independent FRAP events. Scale bars, 2 μm.

We then captured the fusion process of the condensates using time-lapse microscopy. The results show that the condensates formed by all three TFAM proteins can rapidly fuse by necking and relaxation to form a larger one on intersection of two droplets (Figure 5E and Supplemental Video 1-3), suggesting their dynamic property. To validate this, we performed fluorescence recovery after photobleaching (FRAP) analysis to bleach the centers of large droplets and monitored recovery. The bleached pots started to recover after several seconds, and eventually reached around 20% to 50% recovery of the originally detected signal intensity after several minutes (Figure 5F-5K and Supplemental Video 4-6). Together, our findings demonstrated that all three TFAM proteins undergo phase separation *in vitro* and they show varying phase separation property when existed individually.

### TMF selectively recruits TFAMs into phase separated condensates

Given that TMF synergistically acts together with three TFAMs and the four proteins share protein phase separation property, together with the fact that TMF interacts with TFAM1 and TFAM2, we hypothesized that TMF interacts with three TFAMs to form a protein complex that enables formation of a “family condensate”. To test this, we took advantage of the bimolecular fluorescence complementation (BiFC) assay, by which we can simultaneously detect protein-protein interactions and analyze phase-separated condensates in living tomato cells. We performed the pairwise interaction tests between TMF and three TFAMs. The results showed that four proteins interacted with each other in the nucleus, supporting the formation of a protein complex. Image analysis of heterogeneity of the fluorescence intensity and quantification of the cells with condensates indicated that almost all the combinations of interacting pairs induce formation of biological condensates, except the combination of TFAM1 and TFAM2, whose interaction displays homogenous fluorescence signals (Figure 6A, 6B and Supplemental Figure 3). To validate the interactions *in vitro* and monitor the phase separation behavior during interactions without disturbance from other potentially interacting partners, we recombinantly expressed and purified mCherry-TMF (red fluorescence) and GFP-TFAM (green fluorescence) proteins to perform cross-mixing phase separation reactions (Zhou et al., 2020). Apparently, TMF can coexist with itself to form the perfectly merged droplets (Figure 6C). However, it can largely but not fully merge with TFAM1 and TFAM2 droplets (Figure 6C). Surprisingly, TMF shows the most remarkable coincidence with TFAM3 in the same droplets, almost identical to the degree of TMF with itself (Figure 6C), suggesting their tight interactions and cognate property of protein multivalence. In contrast, TMF rarely recruited TFAM11 into its droplets, indicating the selectivity of the recruitment and coexistence during phase separation (Figure 6C). Given the expression of *TFAM11* gradually decreases from early vegetative meristem stage to transitional meristem stage, but dramatically increases at floral meristem stage, apparently opposite to that of *TMF*, the TFAM11 protein might associate with TMF at very early vegetative stages in an unintimate way, and the association was then competed by three other TFAMs due to their closer cognate property of IDR derived multivalences to TMF. Once entering into floral meristem stage, *TMF* disappears, but *TFAM11* up-regulates to take turns to act. Taken together, TMF directly interacts with TFAM1, 2 and 3 and selectively recruits them into phase-separated condensates.

**Figure 6.**
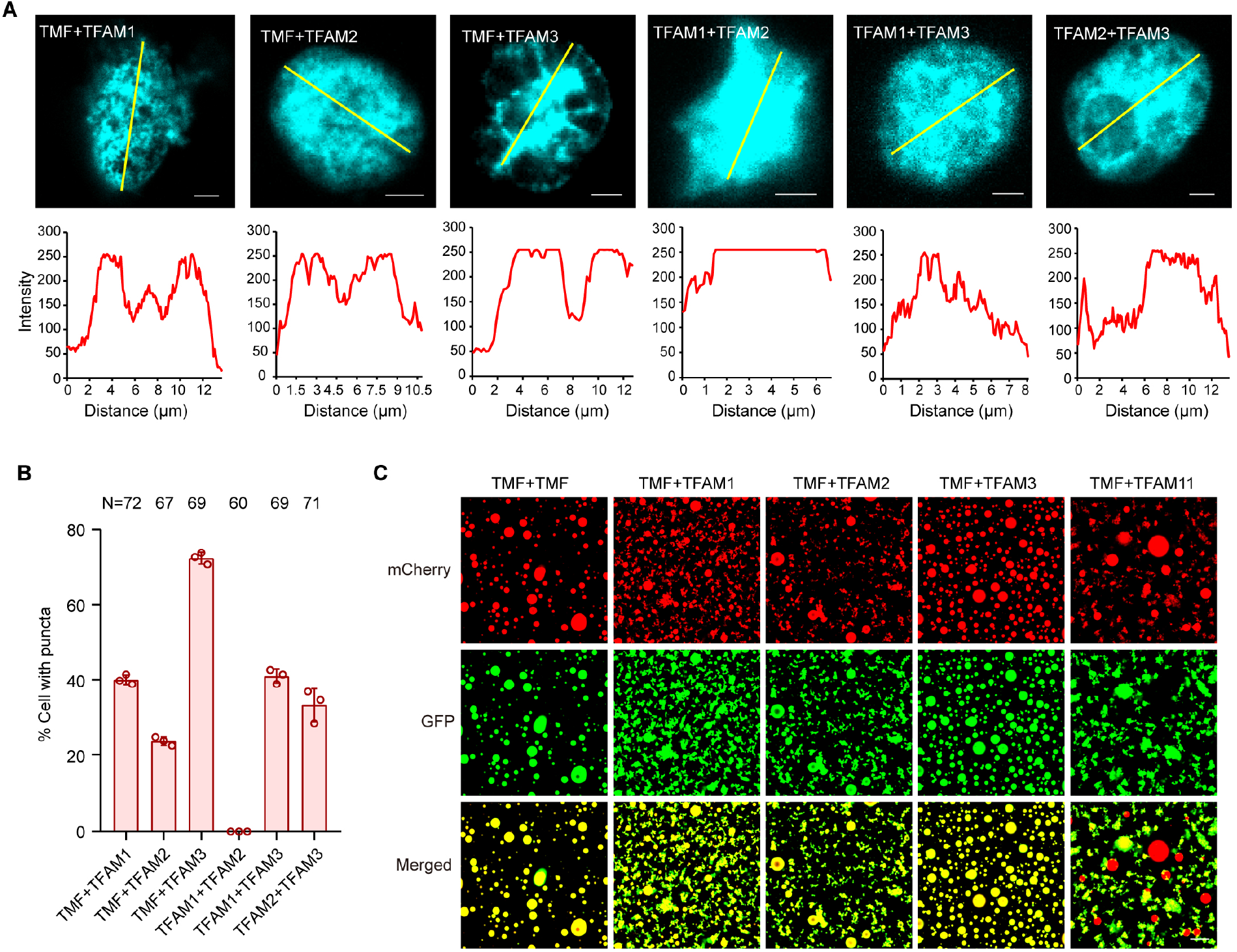
TMF interacts with TFAM proteins to form transcriptional condensates. (**A**) Representative images (upper) showing the interactions between ALOG proteins in BiFC assays. The fluorescence intensity of indicated yellow lines (bottom) showing the heterogenous condensates formed from interactions between TFAM proteins. Scale bars, 2 μm. (**B**) Quantitative data showing the percentage of cells with condensates formed by interactions between TFAM proteins in nuclei. Data are presented as three biological replicates ±SD (n=72, 67, 69, 60, 69, 71). (**C**) Cross-mixing phase separation reactions using recombinantly expressed mCherry-TMF fusion proteins and GFP fusion proteins of TFAMs. Protein concentration, 15 μM; NaCl concentration, 25 mM. Scale bar, 20 μm.

### The ALOG transcriptional condensates repress *AN* expression to synchronize flowering

We recently reported that TMF forms transcriptional condensates to directly target floral identity gene *AN* to repress its expression in the meristems before flowering transition (Huang et al., 2021). To test if TMF acts together with TFAM1/2/3 to target *AN*, we micro-dissected transitional meristems from WT, *tfam1/2/3, tmf* and *tmf tfam1/2* plants for qRT-PCR analysis (Figure 7A left). The results showed that *AN* expression was precociously activated in *tfam1/2/3* comparing to WT (Figure 7A middle). As previously reported and shown here, *AN* prematurely and dramatically upregulated in *tmf* and the effect significantly enhanced in *tmf tfam1/2* triple mutant (Figure 7A right), indicating that TMF and TFAM1/2/3 synergistically repress *AN* expression in the shoot apical meristems at the stages before flowering transition. To validate if the transcriptional repression is direct, we performed a series of transcriptional activity assays using the beta-glucuronidase (GUS)–luciferase (LUC) dual reporter system in tobacco leaves, where promoter sequence of *AN* was fused with GUS to serve as reporter and various combinations of co-expressed TMF and TFAMs served as effectors (Huang et al., 2021) (Figure 7B). The assays showed that coexpression of three TFAMs significantly improved TMF’s repression on the transcription of *AN* (Figure 7C). These results demonstrated that TMF interacts with TFAM1/2/3 to form a transcriptional complex that directly represses *AN* expression.

**Figure 7.**
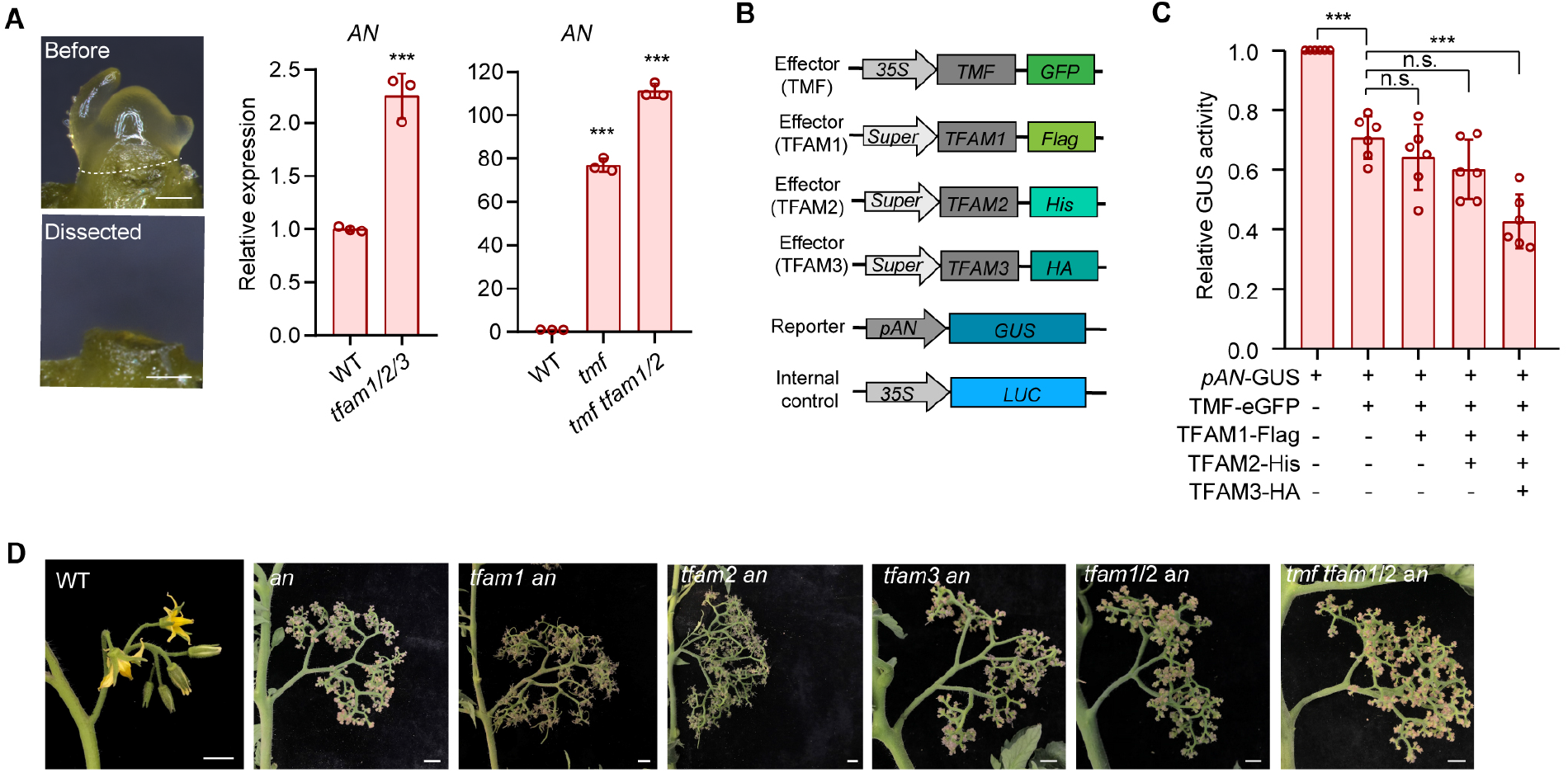
The ALOG transcriptional condensates repress *AN* expression to synchronize flowering. (**A**) Stereoscope images (left) of the micro-dissected transitional meristem for real-time PCR (right). White dashed line indicates the dissection position. The relative expression of *AN* was normalized to WT using *UBIQUITIN* (*UBI*) as an internal control. Data are presented as three repelicates ±SD (n=3, \*\*\**P*<0.001, Student *t*-test). Three independent experiments with similar results were carried out. Scale bars, 100 μm. (**B**) Schematics of constructs used to analyze transcriptional activity. (**C**) Transcriptional repression of *AN* by transcriptional condensates formed from TMF and TFAM proteins. The ratio of GUS to LUC indicates relative transcriptional activity. LUC served as an internal control. Data are presented as six biological replicates from two independent experiments. Data are means ±SD (n=6, \*\*\**P*<0.001, Student *t*-test). (**D**) Representative images for primary inflorescences of WT, *an* and higher-order mutants of *an* and *tfams*. Scale bars, 1 cm.

The aforementioned molecular evidences were then confirmed by extensive genetic analysis. As the *anantha* (*an*) homozygous mutant repeatedly over-proliferates axillary inflorescence meristems but never forms normal flowers (Lippman et al., 2008), we then crossed *anantha* heterozygous mutant with single and high-order mutants of *tfam* and *tmf*. By screening progeny from F2 plants, we obtained *tfam1 an*, *tfam2 an*, *tfam3 an*, *tfam1/2 an* and *tmf tfam1/2 an* mutant. The inflorescences of these mutants were indistinguishable from *anantha* mutant, suggesting that *anantha* mutant is completely epistatic to *tmf* and *tfam* mutants (Figure 7D). Together, these results demonstrate that the transcriptional condensates formed by four paralogous proteins of ALOG family precisely control meristem maturation by directly repressing *AN* expression, which ensures adequate vegetative growth for synchronizing flowering and producing compound inflorescences.

## Discussion

In this study, we uncovered a new mechanism featured by redundant paralogs within one protein family coordinate via phase separation to control a developmental program, by which tomato plants achieve precise control of shoot apical meristem maturation for flowering and compound inflorescence production. Our findings support a model wherein TMF serves a core regulator to selectively recruit its paralogous partners TFAM1, 2 and 3 into phase-separated condensates, the resulting transcriptional condensates bind to the promoter of floral identity gene *AN* to repress it expression during vegetative meristem stages. This phase-separation based repressing program for meristem maturation enables a sufficient duration of vegetative stage for proliferating enough stem cells that ensure synchronization of flowering and production of compound inflorescences (Figure 8). To the best of our knowledge, this is the first example in plants known to elucidate how paralogous transcriptional factors within a protein family synergistically interact to undergo phase separation for stem cell fate decision. The selective-recruitment featured phase separation might represent a general mechanism that is utilized for “task assignment or coordination” among different members within a protein family.

**Figure 8.**
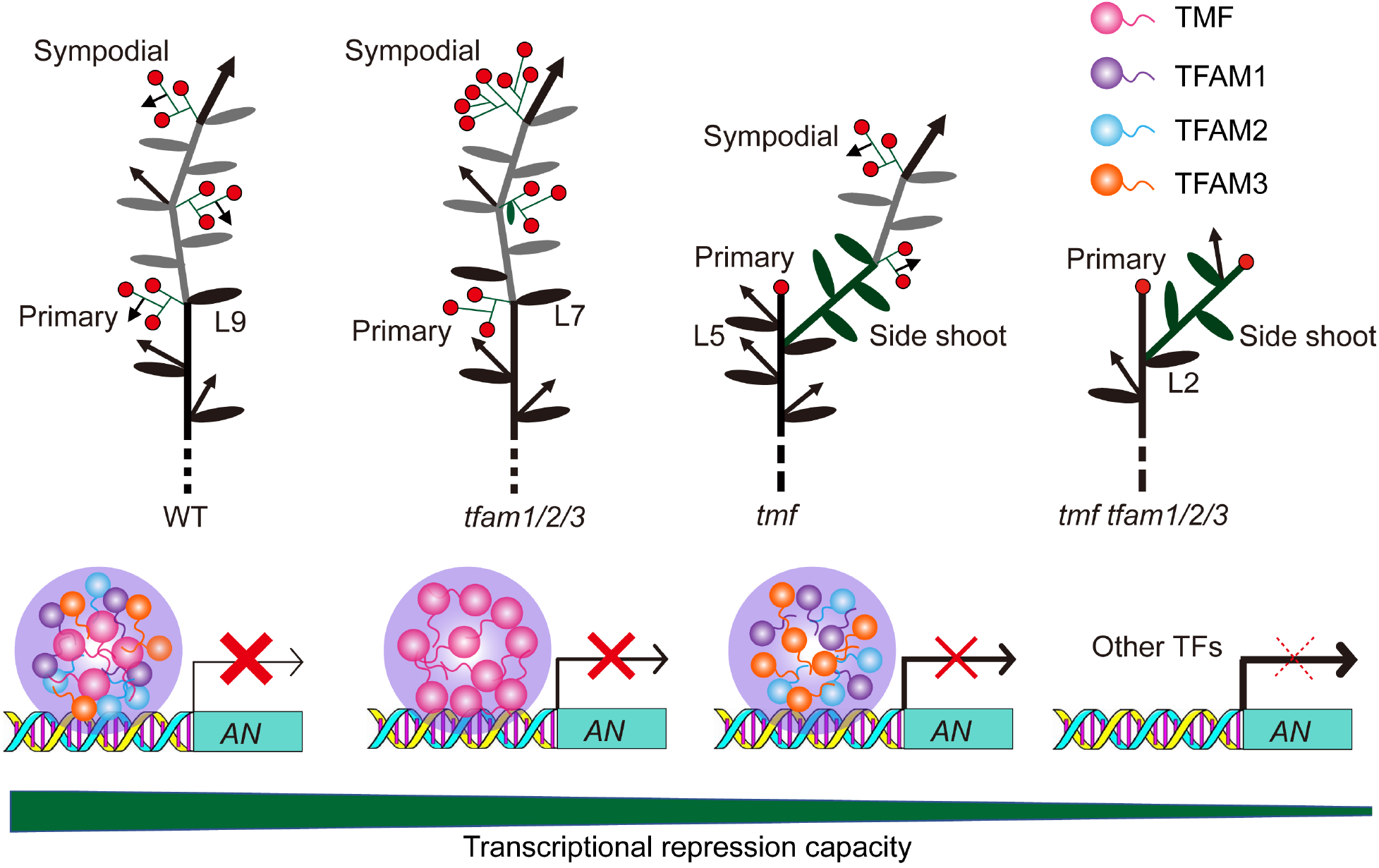
Working model for transcriptional condensates formed by phase-seperated ALOG family proteins in synchronizing flowering and promoting inflorescence complexity.

Despite being sessile, plants have successfully propagated and robustly survived in diverse ecosystems. One key to this evolutionary success is their potent capacity to adapt to local environments (Paaby and Testa, 2018). Biological robustness, usually achieved by the presence of partially redundant parts that result from gene duplication, is an inherent ability of plants that allows them to survive and reproduce in a wide range of different ecosystems (Diss et al., 2014; de Jong and Leyser, 2012). Functional overlap between paralogs allows them to compensate for each other’s loss, as commonly revealed by aggravating genetic interactions (Diss et al., 2014). Creating a full series of mutant combinations of all the ALOG paralogs involved in a certain developmental process allows us to extensively explore the functional overlaps underlying the genetic robustness. For example, when loss of one or two TMF paralogs, tomato plants can still produce multiple-flowered inflorescences with a subtle early-flowering phenotype (Figure 8 and Figure 1D-1F). In this situation, while plants produce fewer fruits and seeds for propagation, their population can still survive. However, once loss of all four functional overlapped ALOG genes simultaneously, tomato plants flower extremely early after producing only two true leaves and develop a single-flowered inflorescence with severe floral organ defects, leading to failure of setting fruits and seeds for propagation (Figure 8 and Figure 2A-2D). Our study discovered that plants adopt a selective-recruitment phase separation mechanism to coordinate the functional overlapped paralogs to achieve biological robustness.

The paralogous proteins within a family often share one or more highly conserved domains that define the protein family, but they often have some regions that are highly diverged. The intrinsically disorder region represents a type of highly-diverged protein sequence. IDRs are featured by their lack of stable secondary or tertiary structure and are rapidly evolving at the primary sequence level. Approximately 40% of all proteins in eukaryotic organisms are either entirely disordered or contain sizeable regions that are disordered (Ward et al., 2004; Zarin et al., 2017). The variance of IDRs among paralogous proteins is likely due to spontaneous mutation occurring during gene duplication (Zarin et al., 2017), which endows the paralogs with different capacity for phase separation and thus varied cognate recognition for functional compensation involved in an essential biological process. IDR driven protein phase separation represents a type of dynamic and flexible protein behavior that has been recently reported to implicate in acclimation responses to cellular pH levels, heat and oxidative stress (Franzmann et al., 2018; Kroschwald et al., 2018; Riback et al., 2017; Yang et al., 2019). The biological condensates formed by functionally associated and phase-separated paralogous proteins might serve as a common mechanism evolved in multicellular organisms for transcriptional adaption to cellular or environmental stresses.

## Methods

### Plant materials and growth conditions

The *tmf, an, tfam1, tfam2* and *tfam3* single mutants used in this study are in tomato (*Solanum lycopersicum*) cultivar M82 background. The higher-order mutants for *tfam1/2*, *tfam1/3*, *tfam2/3*, *tfam1/2/3*, *tmf tfam1*, *tmf tfam2*, *tmf tfam3*, *tmf tfam1/2*, *tmf tfam1/3*, *tmf tfam1/2/3* were produced by crossing using single mutants. The homozygotes were genotyped by digestion of PCR production amplified using the primers listed in Table S1. Seedlings were grown in growth room at 26 °C, with 45–60% relative humidity under LED (Philips Lighting IBRS) light. Greenhouse plants were grown under natural light supplemented with LED. 16 h light/8 h dark photoperiod was used for seedlings and greenhouse plants.

### Transcriptional activity assay

To detect the transcriptional activity of ALOG proteins, the yeast two-hybrid assay was carried out as previously described (Xu et al., 2016). Plasmids for *TMF-BD, TFAM1-BD, TFAM2-BD* were described as previously (Huang et al., 2018) and the coding sequences of *TFAM3* was amplified and cloned into vector pGBKT7 (BD). These plasmids were combined with vector pGADT7 (AD) and co-transformed into AH109 yeast cells, respectively. The transformed cells were plated on the selective mediums.

Meanwhile, the GUS–LUC dual reporter system as previous described was used to perform transcription activity assays *in vivo* (Huang et al., 2021). The ALOG proteins fused different tags served as effctors. *pAN:GUS* served as a reporter, and *35S:LUC* served as an internal control as described previously. Co-infiltrated the plasmids of effector and reporter into *N.benthamiana* leaves, and harvested leaves after 60 h. Total proteins were extracted for measuring the activity of GUS and luciferase (LUC) activity using 4-methylumbelliferyl glucuronide (Sigma) and luciferin (Promega) as substrates, respectively. The transcriptional activity was determined by the ratio of GUS/LUC.

### Recombinant protein expression and purification

To generate the constructs for recombinant protein expression, coding sequences of fusion DNA fragments for *eGFP-TFAMs* and *mCherry-TMF* were cloned into the vector pQE-80L. The plasmids were transformed into *E. coli* Rosetta (DE3) competent cells, and positive bacteria cultured in LB was induced by 0.5 mM isopropyl β-D-1-thiogalactopyranoside (IPTG) for 16 h at 16 °C. Collected cells and performed purification using Ni-NTA (GE healthcare) affinity beads as previous described. Buffer exchange and concentration for eluted proteins were performed using ultrafiltration tubes (Vivaspin turbo). Purified proteins were stored in storage buffer (50 mM Tris-HCl, 200 mM NaCl, pH 7.4) at −80 °C after quick-freezing in liquid nitrogen.

### Phase separation assay and FRAP *in vitro*

The phase separation assays were performed by dilution of purified proteins into buffer containing 50 mM Tris-HCl (pH 7.4) and various concentrations for NaCl to indicated final concentrations in figure legends. Purified proteins were centrifuged 10 min at 14,000g and transferred supernatants into new tubes to exclude the effects caused by precipitated proteins. To generate phase diagram, diluted phase-separated protein solution was incubated for 15 min at room temperature in a 384-well plate. To perform droplet interaction assay for TMF and TFAMs *in vitro*, purified proteins dissolved in buffer containing 50 mM Tris-HCl (pH 7.4), 25 mM NaCl as indicated in figure legends were thoroughly mixed and incubated for 15 min at room temperature in a 384-well plate. Images for droplets and filaments were taken using confocal microscopy (Nikon A1R+) equipped with×20, ×40 and ×100 oil objectives. Fluorescence was excited at 488 and detected at 500–540 for GFP, excited at 543 nm and detected at 595–635 nm for mCherry.

### Subcellular localization and BiFC assays in tomato Protoplasts

To investigate the subcellular localization of TFAM proteins, we generated the constructs. The coding sequences of *TFAM1* and *TFAM3* were amplified and separately cloned into transient expression vector to generate *35S:eGFP-TFAM1*, *35S:TFAM3-eGFP*, and cloned into pSCYNE (SCN) and pSCYCE (SCC) to generate TFAM3-N-CFP (TFAM3-SCN) and TFAM3-C-CFP (TFAM3-SCC) BiFC assay. Plasmids for *35S:TMF-eGFP, 35S:TFAM2-eGFP, TMF-SCC, TMF-SCN, TFAM1-SCC, TFAM1-SCN, TFAM2-SCC* and *TFAM2-SCN* were described as previously (Huang et al., 2018). Plasmids were transfected into protoplasts isolated from tomato cotyledons as previous described. Fluorescent signals detection was performed using confocal microscopy (Leica SP5) with ×20, ×40 objectives.

## Author Contributions

C.X. designed the research; X.H., and N.X. performed most of the experiments and analysed the data; Y.X., L.T., Y.Z., and Y.Y. provided helps in genotyping and plasmid constructions; C.X. wrote the paper with input from X.H..

## Acknowledgments

We thank Z. B. Lippman (Cold Spring Harbor Laboratory) for providing mutant seeds. This study was supported by the Major Research Plan of National Natural Science Foundation of China (31991183), Key Research Program of Frontier Sciences of the Chinese Academy of Science (ZDBS-LY-SM021), the Strategic Priority Research Program of Chinese Academy of Sciences (XDA24030503) to C.X.; China Postdoctoral Science Founding (2018M641516) and a Natural Science Founding (31900174) to X.H.. No conflict interest declared.

## Supplemental information

### Transcriptional condensates formed by phase-separated ALOG family proteins control flowering and inflorescence architecture in tomato

**Supplemental Video 1**. Fusion for two phase-separated GFP-TFAM1 filaments.

**Supplemental Video 2**. Fusion for two phase-separated GFP-TFAM2 filaments.

**Supplemental Video 3**. Fusion for two phase-separated GFP-TFAM3 droplts.

**Supplemental Video 4**. FRAP analysis of phase-separated GFP-TFAM1 *in vitro*.

**Supplemental Video 5**. FRAP analysis of phase-separated GFP-TFAM2 *in vitro*.

**Supplemental Video 6**. FRAP analysis of phase-separated GFP-TFAM3 *in vitro*.

**Figure S1.**
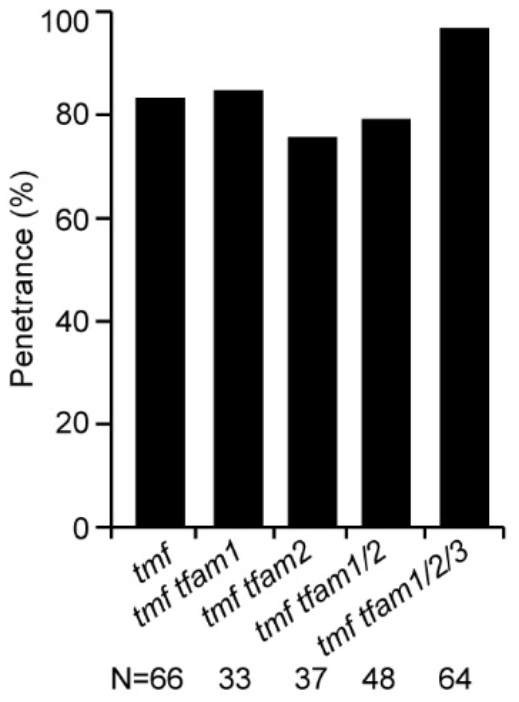
Quantification of penetrance for *tmf* single mutant and higher-order mutants of *tmf* and *tfams*.

**Figure S2.**
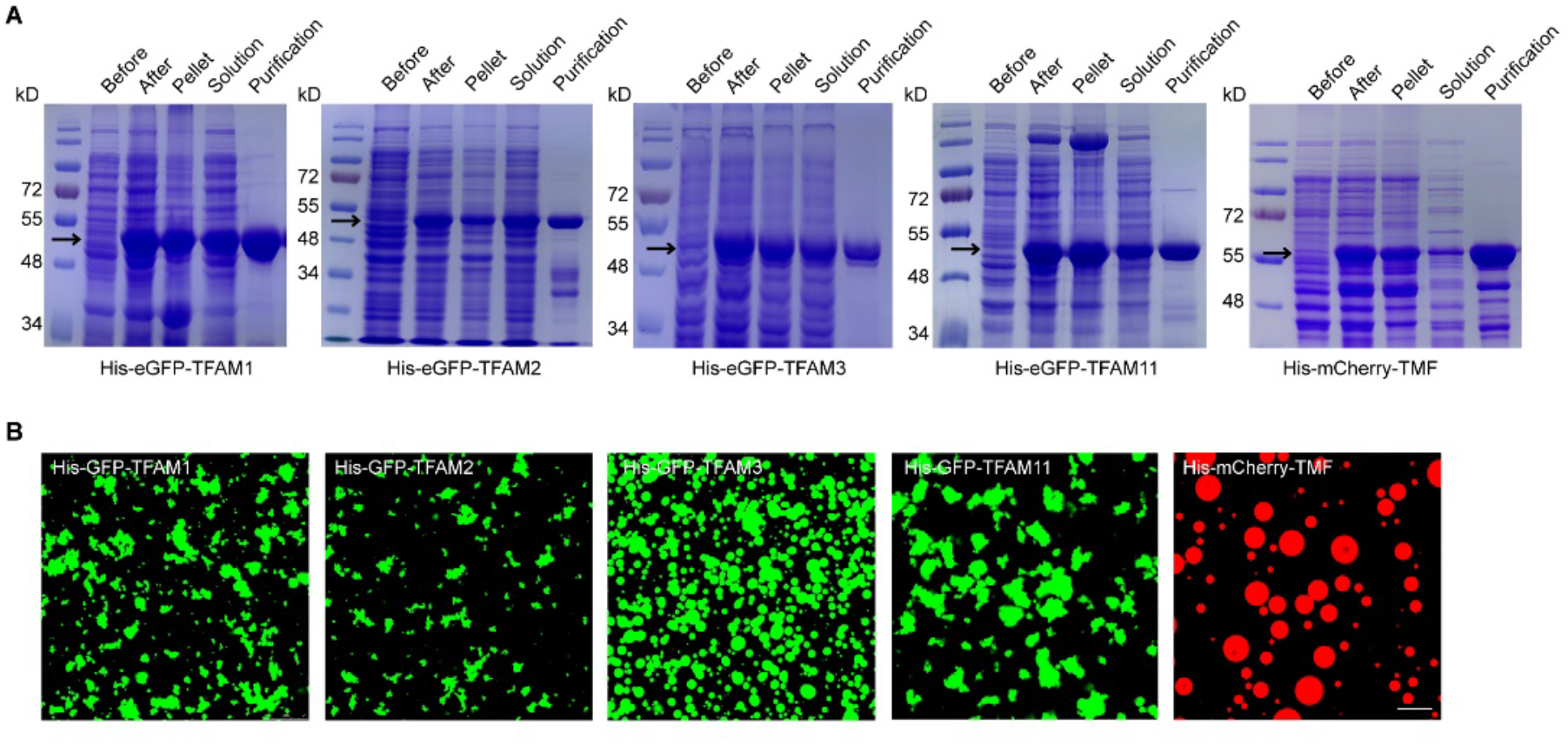
Protein purification for phase separation analysis *in vitro*. (**A**) SDS-PAGE gels showing the inducing and purification for His-eGFP or His-mCherry fused proteins. The black arrows indicate target bands of proteins, respectively. (**B**) Images of phase separation for His-eGFP-TFAMs and His-mCherry-TMF used in this study. Proteins concentration, 15 μM. NaCl concentration, 25 mM. Scale bar, 20 μm.

**Figure S3.**
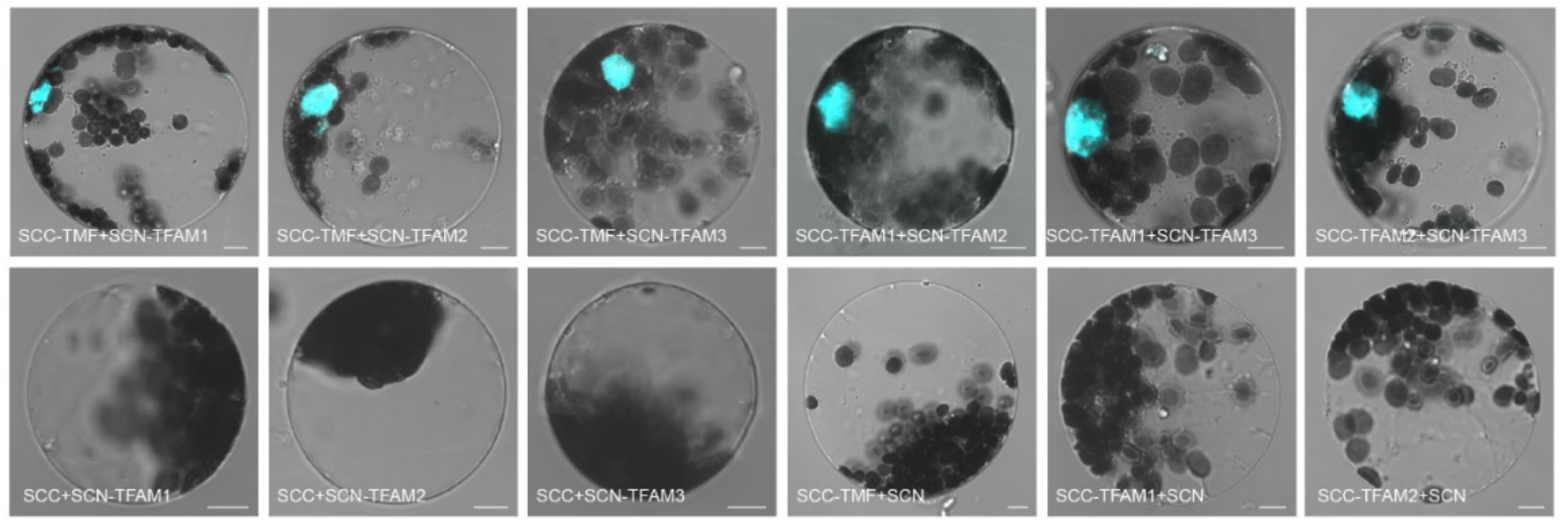
BiFC assays showing the interactions between TMF and TFAMs in nucleus of tomato protoplast. Scale bars, 5 μm.

**Table S1.**
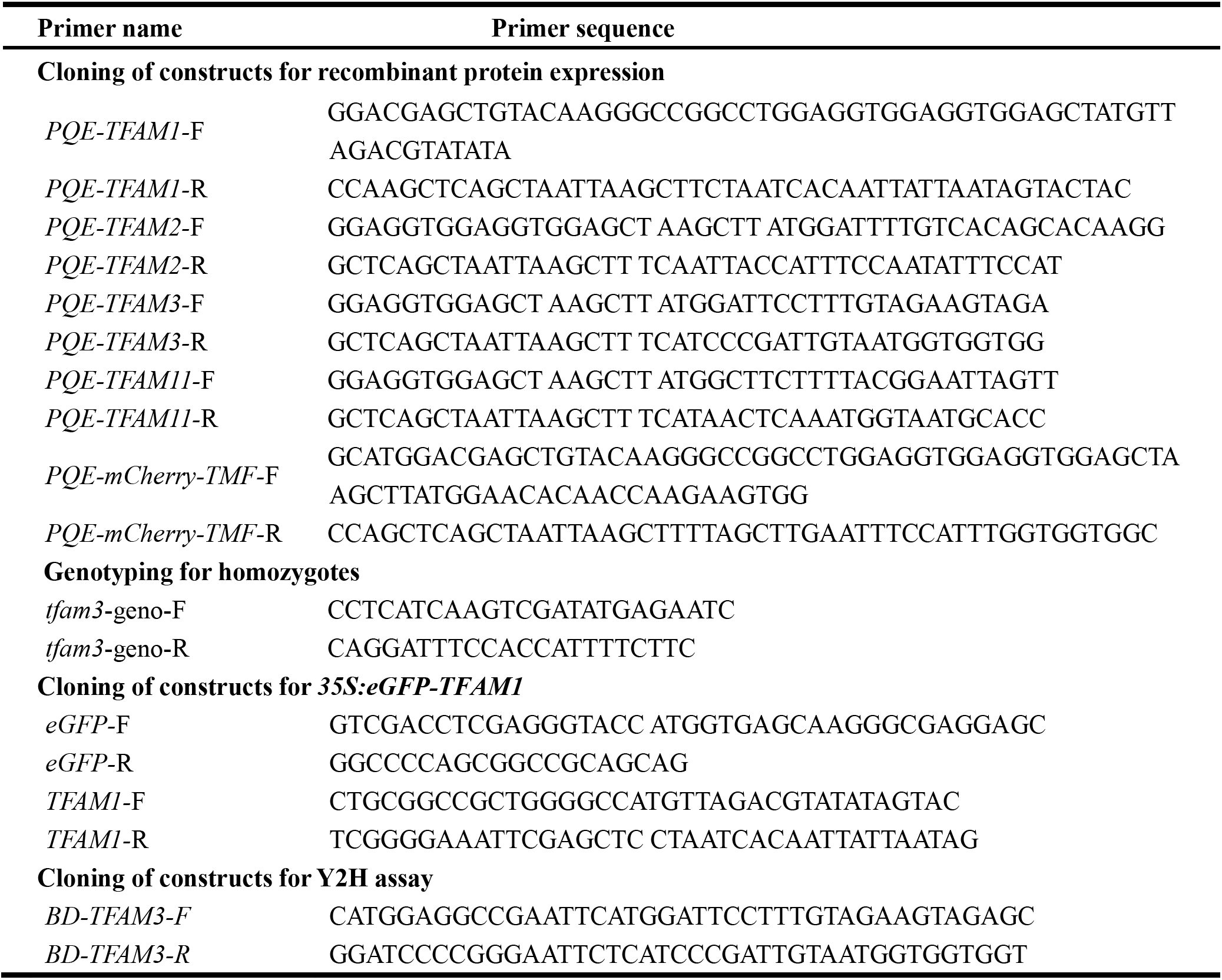
Primer list used in this study

